# LSD1 inhibition attenuates targeted therapy-induced lineage plasticity in *BRAF^V600E^* colorectal cancer

**DOI:** 10.1101/2024.10.25.620306

**Authors:** Christopher A. Ladaika, Averi Chakraborty, Ashiq Masood, Galen Hostetter, Joo Mi Yi, Heather M. O’Hagan

## Abstract

*BRAF* activating mutations occur in approximately 10% of metastatic colorectal cancer (CRCs) and are associated with worse prognosis due to an inferior response to standard chemotherapy. Standard of care for patients with refractory metastatic *BRAF^V600E^* CRC is treatment with BRAF and EGFR inhibitors. However, responses are not durable. Lineage plasticity to neuroendocrine cancer is an emerging mechanism of targeted therapy resistance in several cancer types.

Enteroendocrine cells (EECs), the neuroendocrine cell of the intestine, are uniquely present in *BRAF^V600E^*CRC as compared to *BRAF* wildtype CRC. Here, we demonstrated that combined BRAF and EGFR inhibition enriches for EECs in several models of *BRAF^V600E^*CRC. Additionally, EECs and other secretory cell types were enriched in a subset of *BRAF^V600E^* CRC patient samples following targeted therapy. Importantly, inhibition of the lysine demethylase LSD1 with a clinically relevant inhibitor attenuated targeted therapy-induced EEC enrichment through blocking the interaction of LSD1, CoREST2 and STAT3.

**Statement of Significance:** Our findings that BRAF plus EGFR inhibition induces lineage plasticity in *BRAF^V600E^* CRC represents a new paradigm for how resistance to BRAF plus EGFR inhibition occurs and our finding that LSD1 inhibition blocks lineage plasticity has the potential to improve responses to BRAF plus EGFR inhibitor therapy in patients.

## Introduction

*BRAF* activating mutations occur in approximately 10% of metastatic colorectal cancers (CRCs) and are associated with poor prognosis due to an inferior response to high-dose 5-Flurouracil- based chemotherapy (1). *BRAF^V600E^* mutation activates mitogen-activate protein kinase (MAPK) signaling, a pathway that has been a significant focus for targeted therapy approaches. Findings from the BEACON (Binimetinib, Encorafenib, And Cetuximab cOmbiNed) clinical trial resulted in the standard of care for patients with refractory *BRAF^V600E^* metastatic CRC becoming combined BRAF (encorafenib) and epidermal growth factor receptor (EGFR; cetuximab) inhibition (2).

However, the median overall survival of patients receiving combined therapy is only 9 months.

Targeted therapies have been developed for many cancers to inactivate mutated or hyperactive proteins that drive cancer-promoting signaling pathways. However, cancers have developed methods of resistance to recent potent targeted therapies that include cancers undergoing shifts in cell identity resulting in cancer cells that are no longer reliant on the signaling pathway being targeted (3). For example, transformation from adenocarcinoma to neuroendocrine prostate cancer is a mechanism of resistance to treatment with androgen receptor signaling inhibitors in about 20% of castrate-resistant prostate cancer (4). Additionally, treatment of non-small cell lung cancer (NSCLC) with EGFRi induces transdifferentiation to SCLC, which has a poorly differentiated neuroendocrine phenotype that is correlated with rapid growth and a high metastatic rate (5).

The normal intestinal epithelium contains multiple cell types, including absorptive enterocytes and secretory goblet and enteroendocrine cells (EECs) (6). Factors released by secretory cells have roles in the normal colon that promote wound healing, gut integrity, and immune cell regulation but can also be tumor promoting (7). Furthermore, secretory progenitors and EECs are capable of reverting to stem cells after crypt damage (8,9), which gives these cells the capability of promoting tumor survival under stressful conditions such as during metastasis or treatment. We have demonstrated that EECs and goblet cells are enriched in *BRAF^V600E^* CRC and promote cancer cell survival via secreted factors (10). LSD1 (KDM1A), a lysine demethylase overexpressed in CRC, has been implicated in fate specification of multiple cellular lineages in normal and cancerous cells (11). We have demonstrated that LSD1 is a major regulator of EEC differentiation in *BRAF^V600E^* CRC through interacting with CoREST2 and potentiating STAT3 activity (10,12).

In the normal colon, MAPK pathway inhibition pushes secretory cell differentiation toward the EEC lineage (13). Therefore, we sought to determine how standard of care treatment for refractory metastatic *BRAF^V600E^* CRC, which targets the MAPK pathway, alters cancer epithelial cell types in *BRAF^V600E^*CRC. We determined that combined BRAFi plus EGFRi specifically enriched for EECs in several cell line, organoid and colon orthotopic in vivo models of *BRAF^V600E^*CRC. Furthermore, we demonstrated that LSD1 promoted tumor heterogeneity and lineage plasticity in *BRAF^V600E^* CRC and that targeting LSD1 by epigenetic therapy improved responses to BRAFi plus EGFRi by blocking therapy-induced lineage plasticity.

## Results

### Combined BRAF plus EGFR inhibition enriches for EECs in *BRAF^V600E^* CRC

As targeted therapies have been shown to induce cell plasticity in other cancer types, we treated *BRAF^V600E^* CRC HT29 and NCI-H508 cell lines with BRAFi (encorafenib) and EGFRi (gefitinib) alone or in combination and assayed changes in expression of marker genes for different colon epithelial cell types. In HT29 cells, BRAFi alone or in combination with EGFRi increased expression of EEC marker genes *NGN3* and *INSM1*, but not marker genes of goblet cells (*MUC2*) or enterocytes (*HES1*) (Fig. 1A). As EEC marker gene expression increased with length of treatment and doses of encorafenib between 1 and 100 nM had similar effects on expression of EEC genes, most future experiments were performed with 4 days of treatment at 2.5 nM encorafenib (Fig. S1A, S1B). In NCI-H508, cells EGFRi alone or in combination with BRAFi increased expression of EEC marker genes (Fig. 1B). Consistent with the gene expression findings, in HT29 cells, BRAFi or BRAFi+EGFRi increased the percentage of cells positive for β3-tubulin, a marker of neuronal cells that is also expressed in EECs (Fig. 1C). In NCI-H508 cells, EGFRi or BRAFi+EGFRi increased the percentage of β3-tubulin positive cells (Fig. 1D). BRAFi+EGFRi also increased expression of EEC marker genes *NGN3* and *INSM1* in a *BRAF^V600E^*CRC human organoid model 817 without changing expression of goblet or enterocyte markers (Fig. 1E). As MEK inhibitors (MEKi) have also been used to treat patients with metastatic *BRAF^V600E^* CRC, we tested the effect of MEKi on marker gene expression in HT29 cells. Increasing concentrations of MEKi alone increased expression of EEC genes *NGN3* and *INSM1* and goblet gene *MUC2* and, to a lesser extent, enterocyte gene *HES1* (Fig. S1C). MEKi alone or in combination with EGFRi also increased the percentage of β3-tubulin positive cells (Fig. S1D).

**Figure 1.**
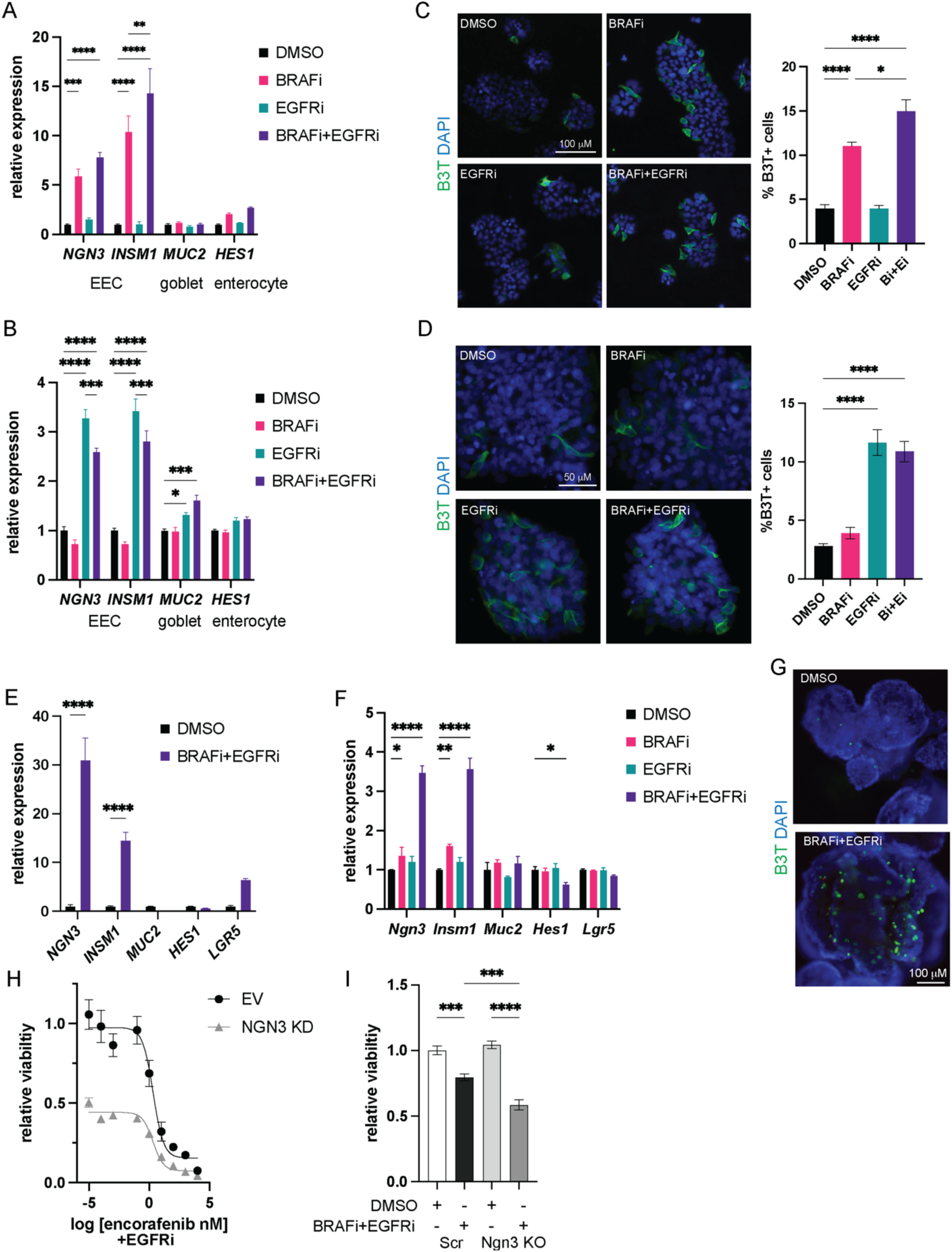
BRAF plus EGFR inhibition enriches for EECs in *BRAF^V600E^* CRC. (A) Relative gene expression of indicated genes in HT29 cells treated with DMSO or 2.5 nM encorafenib (BRAFi) alone or in combination with 500 nM gefitinib (EGFRi) for 48H. Gene expression was normalized to the housekeeping gene *RhoA* and then to the DMSO treated cells. Graph represents mean +/- SEM. N=3. (B) Relative gene expression of indicated genes in NCI-H508 cells treated with DMSO or 2.5 nM encorafenib alone or in combination with 250 nM gefitinib for 48H. Data is normalized and presented as in A. (C) Immunofluorescence for β3-tubulin (B3T) in HT29 cells treated as in A for 72H. Graph is the %β3-tubulin+ cells of the total number of cells per field. N=3. (D) Immunofluorescence for β3-tubulin (B3T) in NCI-H508 cells treated as in B for 72H and analyzed as in C. (E) Relative gene expression of indicated genes in 817 human CRC organoids treated with DMSO or 2.5 nM encorafenib and 500 nM gefitinib for 6 days. Data is normalized and presented as in A. (F) Relative gene expression of indicated genes in TP KO mouse CRC organoids treated with DMSO or 2.5 nM encorafenib alone or in combination with 500 nM gefitinib for 4 days. Data is normalized and presented as in A. (G) Immunofluorescence for β3-tubulin (B3T) in TP KO organoids treated as in F. (H) Encorafenib dose response curve of empty vector (EV) or NGN3 knockdown (KD) HT29 cells treated with 500 nM gefitinib (EGFRi) for 72H. Viability was normalized to DMSO treated EV cells. N=3. (I) Relative viability of TP KO scramble (Scr) and NGN3 KO organoids treated as in F. Viability was normalized to DMSO treated scramble cells. N=3. Significance was determined by one-way ANOVA with Tukey pairwise multiple comparison testing. *P≤ 0.05, ** P ≤ 0.01, *** P ≤ 0.001, **** P ≤ 0.0001

To generate a model of metastatic *BRAF^V600E^* CRC to use in vitro and in immunocompetent mice, we derived tumor organoids from tumors from an in situ model of *BRAF^V600E^Apc^λ1716/+^* tumorigenesis (14,15). Because we were interested in ultimately studying metastatic CRC, we then knocked out Tgfbr2, which has reduced expression in *BRAF* mutant CRC as compared to normal colon and *BRAF* wildtype CRC, and Trp53 in the organoids using a CRISPR/Cas9 approach (Fig. S1E,S1F). Interestingly, the Tgbfr2 + Trp53 knockout (TP KO) organoids had reduced expression of Trp53 target gene *Bax1*, and tumor suppressor gene *Rb1,* as well as increased expression of EEC marker genes, *Ngn3* and *Insm1*, and stem cell gene *Lgr5* relative to the mock KO organoids (Fig. S1G, S1H). Single agent treatment only had modest effects on expression of marker genes in the TP KO organoids, whereas combined BRAFi+EGFRi treatment significantly increased EEC marker gene expression (Fig. 1F, S1I). Immunofluorescence for β3-tubulin confirmed that BRAFi+EGFRi treatment increased the percentage of EECs in the organoids (Fig. 1G).

To determine if the presence of EECs influences the sensitivity of the cells to treatment, we knocked down NGN3 in HT29 cells (Fig. S1J) and performed an encorafenib dose response curve in the presence of a constant concentration of gefitinib. NGN3 knockdown (KD) cells had decreased viability relative to empty vector (EV) control KD cells and were more sensitive to encorafenib treatment (Fig. 1H). We also knocked out Ngn3 in the TP KO organoids to reduce EECs (Fig. S1K). While Ngn3 KO had no effect on the baseline viability of the TP KO organoids, BRAFi+EGFRi reduced viability of the Ngn3 KO organoids to a greater extent than the scramble control organoids (Fig. 1I). Altogether, these findings demonstrate that BRAFi+EGFRi treatment enriches for EECs in *BRAF^V600E^* CRC and suggests that EECs promote resistance to this therapy.

### Residual tumors following BRAF plus EGFR inhibitor treatment have altered tumor cell composition

To determine how BRAFi+EGFRi treatment alters tumor composition in vivo, we orthotopically engrafted HT29 cells expressing tdTomato and luciferase or 817 organoids into the colons of NSG mice (Fig. S2A). After tumor formation, mice were treated for 3 weeks with encorafenib (BRAFi) and cetuximab (EGFRi). The treatment prevented and reduced growth of HT29 and 817 orthotopic tumors, respectively (Fig. 2A, 2B) and significantly reduced lung metastasis of both tumor types (Fig. S2B). Treatment reduced levels of phosphorylated ERK (pERK) as expected and reduced Ki67 staining without increasing the apoptosis marker, cleaved caspase 3 (Fig. S2C, S2D). Interestingly, by H&E staining, all residual tumors had increased intratumoral cystic spaces (Fig 2C, 2D). Consistent with our in vitro data, the residual HT29 tumors from treated mice were enriched for EECs as indicated by increased synaptophysin, β3-tubulin, and INSM1 staining (Fig. 2C, 2D, S2E). Synaptophysin β3-tubulin and INSM1 positive cells were also increased in 817 tumors from BRAFi+EGFRi compared to vehicle treated mice but, unlike in the HT29 tumors, the positive cells were limited to small pockets (Fig. 2E).

**Figure 2.**
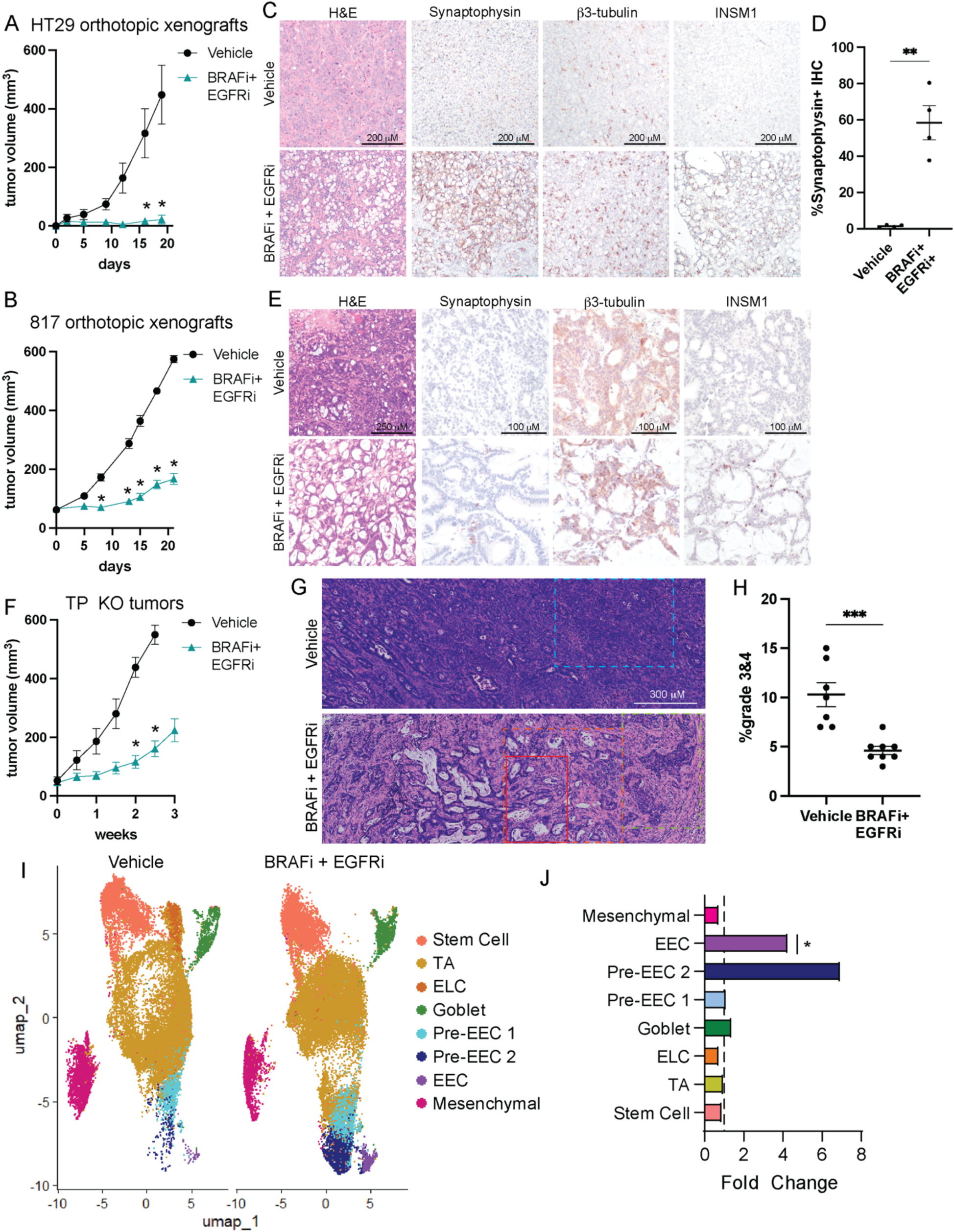
BRAFi plus EGFRi treatment induces a neuroendocrine phenotype in residual *BRAF^V600E^* tumors. (A) Following tumor formation of HT29 cells or (B) 817 human CRC organoids orthotopically engrafted into the colons of NSG mice, mice were treated with vehicle or BRAFi (encorafenib, 20 mg kg^-1^, daily) and EGFRi (cetuximab, 20 mg kg^-1^, biweekly). Graph is the mean +/- SEM of the tumor volume measured by caliper over the course of treatment. N=5 mice per group. (C) Representative H&E and IHC images of HT29 colon tumors from the experiment in A. (D) IHC scoring of synaptophysin staining. Each point represents the scoring for one tumors. Lines represent mean -/+ SEM. (E) Representative H&E and IHC images of 817 colon tumors from the experiment in B. (F) Following tumor formation of TP KO mouse CRC organoids orthotopically engrafted into the colons of C57Bl/6 mice, mice were treated with vehicle or BRAFi (encorafenib, 20 mg kg^-1^, daily) and EGFRi (gefitinib, 75 mg kg^-1^, daily). Tumor growth was measured and plotted as in A. N=7, vehicle. N=8, BRAFi+EGFRi. (G) Representative H&E images of TP KO tumors. Blue dashed box indicates undifferentiated tumor glands, grade 3,4 solid and cord like tumor with dense stroma. Red solid box indicates well differentiated glandular tumor with goblet cells and active mucin production. Orange dashed box indicates well defined mature stroma. Green dashed box indicates less differentiated solid tumor area with biphasic appearance suggestive of neuroendocrine features. (H) Semi quantitative percentage of TP KO tumor areas that are grade 3&4. Each point represents the scoring for one tumor. Lines represent mean -/+ SEM. (I) UMAP dot plot of TP KO tumor epithelial cell scRNAseq data from mice treated with vehicle (left) or BRAFi+EGFRi (right). Samples are colored by cell type/cluster. TAs are transit amplifying cells, ELCs are enterocyte-like cells, and EECs are enteroendocrine cells. (J) Fold change in cell type proportions in BRAFi+EGFRi relative to vehicle samples. Significance was determined by two-way ANOVA corrected for multiple comparisons using the Šídák method (A,B,F), student’s T-test (D,H) and by a moderated t-test (J). *P≤ 0.05.

To explore how BRAFi+EGFRi therapy alters tumor composition in a syngeneic model of *BRAF^V600E^* CRC, we implanted TP KO organoids expressing luciferase into the colons of wildtype C57Bl/6 mice (Fig. S2F). As cetuximab is an anti-human EGFR antibody it lacks activity on mouse EGFR. Therefore, we treated tumor bearing C57Bl/6 mice with encorafenib and the EGFR tyrosine kinase inhibitor gefitinib daily for 3 weeks. Treatment reduced tumor growth and metastasis to the lungs and iliac and lumbar lymph nodes (Fig. 2F, S2G, S2H).

Microscopic review revealed that all orthotopic tumors contained regions of mixed histology by tumor grade. The vehicle tumors contained more solid and poorly differentiated regions in contrast to the BRAFi+EGFRi tumors (Fig. 2G, 2H). In addition, the treated tumors showed significantly increased areas of differentiated glandular structures and were intimately associated with more mature stroma characterized by more ordered, fibrillar and eosinophilia compared to the vehicle tumors (Fig. 2G). To further explore how therapy alters tumor cell composition, we performed FLEX single cell RNA sequencing (scRNAseq) on 7 vehicle and 8 BRAFi+EGFRi tumors. To focus on tumor epithelial cell populations, clustering analysis was performed using only tumor epithelial cells extracted from the scRNAseq data. Clusters were defined by expression of marker genes, cell cycle profile and trajectory analysis (Fig. S2I, S2J, Supplementary Table S1). Furthermore, cluster analysis of module scores for Hallmark gene sets confirmed cluster identification, with stem cells being enriched for MYC and E2F targets, mesenchymal cells being enriched for epithelial to mesenchymal transition (EMT) and TGF beta signaling pathways and EECs being enriched for pancreas beta cells, a cell type that shares many gene expression pathways with EECs. Examining differences in cell type proportions demonstrated significant enrichment of differentiated EECs and a trend towards enrichment of Pre-EEC 2 cells in BRAFi+EGFRi tumors as compared to vehicle tumors (Fig. 2J). Interestingly, *Prox1*, a gene expressed by progenitor cells and EECs in the normal intestine and related to neuroendocrine plasticity in prostate cancer (9,16), was identified as a marker gene of all pre- EEC and EEC clusters (Fig. S2L). Regions of PROX1+ cells were present in all tumors regardless of treatment with some of the highest positivity occurring in BRAFi+EGFRi tumors (Fig. S2M). PROX1+ cells were also present in HT29 orthotopic tumors and increased in the HT29 BRAFi+EGFRi tumors relative to the vehicle tumors (Fig. S2N).

### Inhibition of LSD1 reduces BRAFi+EGFRi-induced cell plasticity

Because we had previously demonstrated that LSD1 promotes EEC differentiation in *BRAF^V600E^*CRC (10,12), we sought to determine if LSD1 inhibition also promotes therapy-induced EEC enrichment. LSD1 knockdown or inhibition with the LSD1/CoREST dual inhibitor corin reduced the BRAFi-induced increase in expression of EEC marker genes in HT29 cells ((17); Fig. S3A, S3B). LSD1 knockdown or inhibition with corin also increased the sensitivity of HT29 cells to encorafenib treatment (Fig. S3C, S3D).To identify a clinically relevant inhibitor to use in our studies, we tested the effect of a panel of LSD1 inhibitors on EEC differentiation (11). Only SP- 2577 reduced the basal expression of EEC marker genes in HT29 cells even though all the inhibitors reduced global levels of H3K4me2 (Fig. S3E, S3F). SP-2577 also reduced the BRAFi+EGFRi-induced increase in EEC marker gene expression (Fig. 3A, 3B, S3G) and increase in β3-tubulin+ EECs (Fig. 3C, 3D) in HT29 and H508 cells. Similarly to LSD1 knockdown, SP-2577 increased the sensitivity of HT29 cells to encorafenib treatment and the sensitivity of HT29 and NCI-H508 cells to combined BRAFi+EGFRi treatment (Fig. S3H, 3E, 3F). SP-2577 treatment also blocked the MEKi-induced increase in expression of EEC marker genes, the MEKi+EGFRi-induced increase in β3-tubulin+ EECs and increased the sensitivity of HT29 cells to MEKi (Fig. S3I, S3J, S3K).

**Figure 3.**
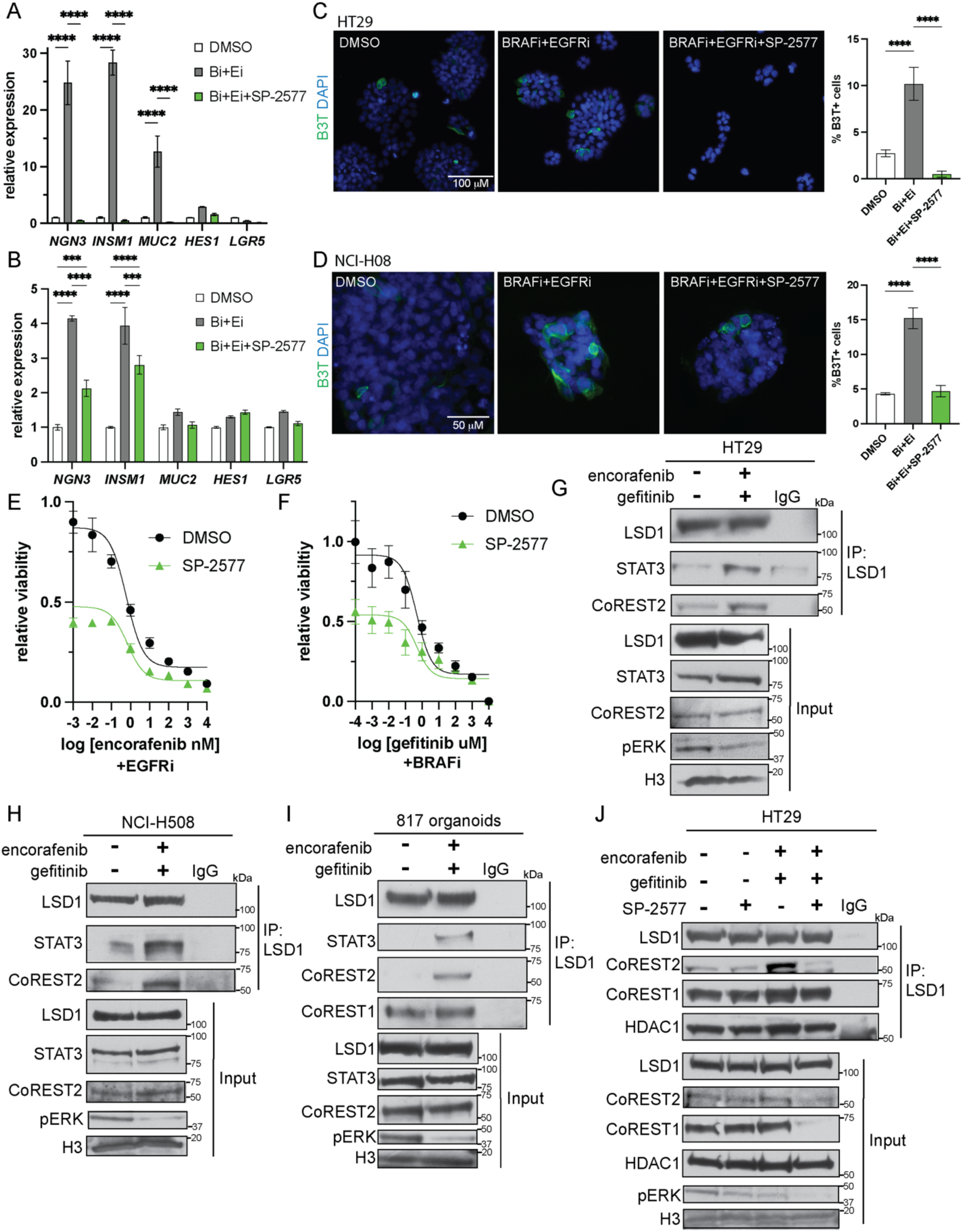
LSD1 inhibition blocks therapy-induced EEC enrichment in *BRAF^V600E^* CRC. (A) Relative gene expression of indicated genes in HT29 cells treated with DMSO or 2.5 nM encorafenib (BRAFi, Bi) plus 500 nM gefitinib (EGFRi, Ei) with or without 500 nM SP-2577 (LSD1i) for 4 days. Gene expression was normalized to the housekeeping gene RhoA and then to the DMSO treated cells. Graph represents mean +/- SEM. N=3. (B) Relative gene expression of indicated genes in NCI-H508 cells treated with DMSO or 2.5 nM encorafenib plus 250 nM gefitinib for with or without 500 nM SP-2577 for 48H. Data is normalized and presented as in A. (C) Immunofluorescence for β3-tubulin (B3T) in HT29 cells treated as in A for 72H. Graph is the %β3-tubulin+ cells of the total number of cells per field. N=3. (D) Immunofluorescence for β3- tubulin in NCI-H508 cells treated as in B and analyzed as in C for 72H. (E) Encorafenib dose response curve of HT29 cells treated with 500 nM gefitinib (EGFRi) and DMSO or 500 nM SP- 2577 for 72H. Viability was normalized to cells treated with DMSO only. (F) Gefitinib dose response curve of NCI-H508 cells treated with 2.5 nM encorafenib (BRAFi) and DMSO or 500 nM SP-2577 for 72H. Normalized as in E. (G) LSD1 coIP in nuclear lysates prepared from HT29 cells treated with DMSO or 2.5 nM encorafenib plus 500 nM gefitinib. (H) LSD1 coIP in nuclear lysates prepared from NCI-H508 cells treated with DMSO or 2.5 nM encorafenib plus 250 nM gefitinib. (I) LSD1 coIP in nuclear lysates prepared from 817 organoids treated with DMSO or 2.5 nM encorafenib plus 500 nM gefitinib. (J) LSD1 coIP in nuclear lysates prepared from HT29 cells treated as in A. Significance was determined by one-way ANOVA with Tukey pairwise multiple comparison testing. *P≤ 0.05, ** P ≤ 0.01, *** P ≤ 0.001, **** P ≤ 0.0001.

We recently determined that LSD1 in combination with CoREST2 promotes EEC differentiation by potentiating STAT3 activity (12). Consistent with our previous findings, STAT3i reduced the baseline and BRAFi-induced expression of EEC markers in HT29 cells (Fig. S3L).

BRAFi+EGFRi treatment induced the interaction of LSD1 with STAT3 and CoREST2 in HT29 and H508 cells and 817 organoids (Fig. 3G, 3H, 3I). Interestingly, combing SP-2577 with BRAFi+EGFRi treatment blocked the inhibitor-induced increased interaction between LSD1 and CoREST2, without effecting the interaction between LSD1 and CoREST1 or HDAC1 (Fig. 3J). SP-2577 also reduced the inhibitor-induced interaction between LSD1 and CoREST2 and STAT3 in NCI-H508 cells (Fig. S3M). Altogether this data, demonstrates that LSD1 is critical for BRAFi+EGFRi-induced EEC differentiation and that SP-2577 blocks BRAFi+EGFRi-induced EEC differentiation likely by disrupting the LSD1-CoREST2-STAT3 interaction.

### LSD1 inhibition attenuates therapy-induced cell plasticity in vivo

We next explored connections between LSD1-CoREST2-STAT3 and EECs in our in vivo models of *BRAF^V600E^* CRC. In our scRNAseq data from TP KO colon orthotopic tumors, *Rcor2*, the gene encoding CoREST2, was predominantly expressed in the Pre-EEC 2 and EEC clusters in BRAFi+EGFRi tumors (Fig. 4A). Additionally, gene set module score analysis demonstrated that EECs were enriched for genes that were down-regulated after LSD1 knockout in the pituitary relative to the other cell populations and for genes that are regulated by repressor element-1 silencing transcription factor/neuron-restrictive silencer factor (REST/NSRF), a repressor of neuroendocrine genes (Fig. 4B, 4C) (18). These computational analyses suggest that LSD1-CoREST2 contributes to EEC differentiation in *BRAF^V600E^* CRC following BRAFi+EGFRi treatment in vivo. To further confirm these findings, we performed LSD1 and STAT3 co-IPs from nuclear lysates prepared from HT29 and 817 orthotopic tumors from treated mice. The interaction between LSD1 and STAT3 and CoREST2 and STAT3 and LSD1 was higher in BRAFi+EGFRi than vehicle tumors (Fig. 4D, S4A, S4B).

**Figure 4.**
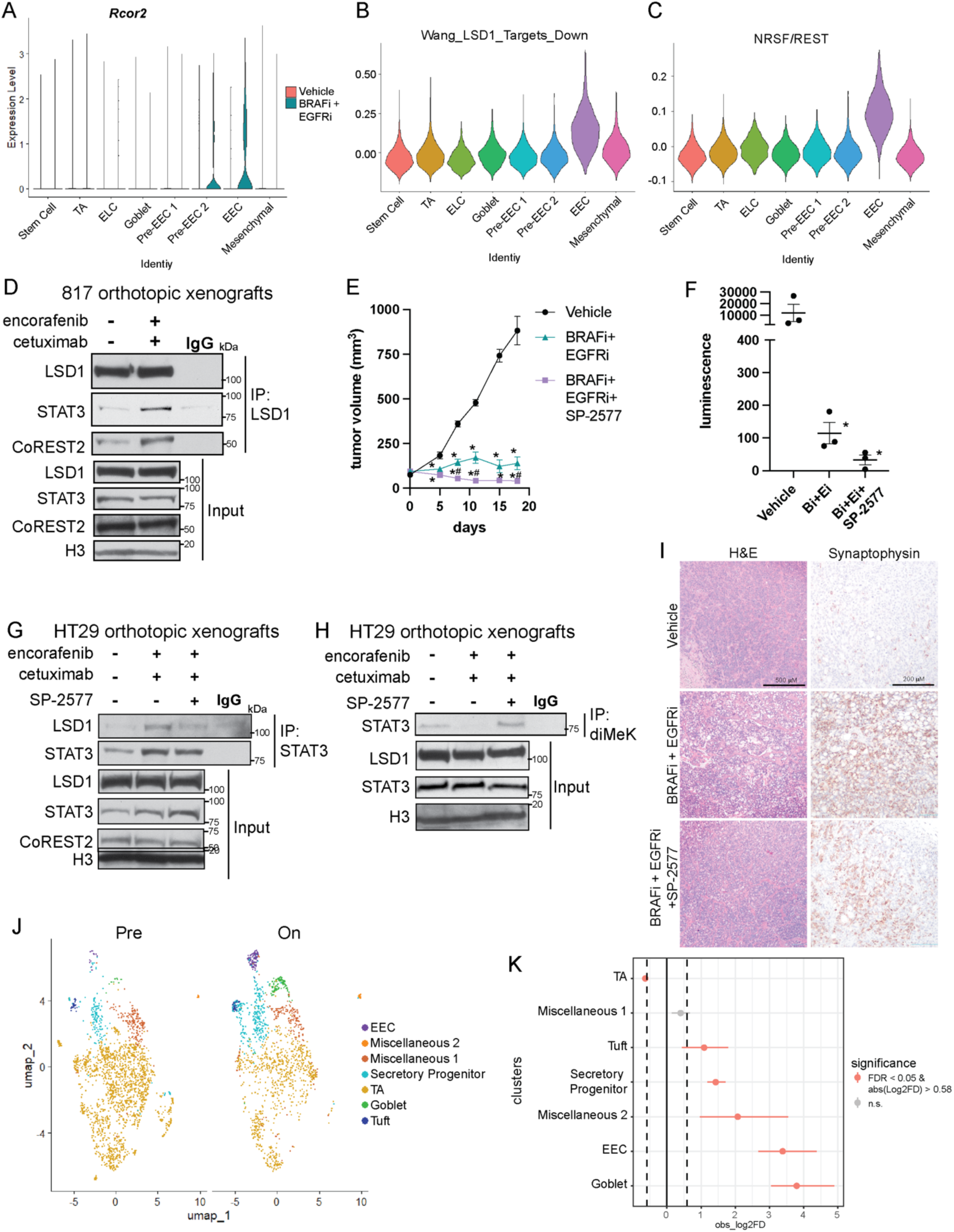
LSD1 inhibition blocks BRAFi+EGFRi-induced lineage plasticity in vivo. (A) Rcor2 violin plot from TP KO scRNAseq data. (B) Module scores for the Wang LSD1 Targets Down gene set for each epithelial cell in the TP KO scRNAseq data. (C) Module scores for the NRSF gene set for each epithelial cell in the TP KO scRNAseq data. (D) LSD1 coIP in nuclear lysates prepared from 817 colon orthotopic tumors from vehicle or encorafenib + cetuximab treated NSG mice. (E) Following tumor formation of HT29 cells orthotopically engrafted into the colons of NSG mice, mice were treated with vehicle or BRAFi (encorafenib, 20 mg kg^-1^, daily) and EGFRi (cetuximab, 20 mg kg^-1^, biweekly) with or without SP-2577 (LSD1i, 100 mg kg^-1^, twice daily). Graph is the mean +/- SEM of the tumor volume measured by caliper over the course of treatment. N=5 mice per group at days 0-7. N=3 mice per group at days 12-18. *P≤ 0.05 relative to vehicle. ^#^P≤ 0.05 relative to BRAFi+EGFRi. Significance was determined by fitting a mixed model and correcting for multiple comparisons by controlling the false discovery rate using the two-stage step-up method of Benjamini, Krieger and Yekutieli. (F) Total luminescence signal detected by plate reader from lungs incubated ex vivo in PBS+luciferin. A lung from a non-tumor bearing mouse was used as a negative control. Each point represents an individual lung. Lines represent mean -/+ SEM. *P≤ 0.05 relative to vehicle. Significance was determined by one-way ANOVA with Tukey pairwise multiple comparison testing. (G) STAT3 coIP in nuclear lysates prepared from HT29 colon orthotopic tumors from mice treated with vehicle, encorafenib + cetuximab, or encorafenib + cetuximab + SP-2577 for 7 days. The same tumor lysates in the left panel were also used in the experiment in S4A so the input blots are the same. (H) Dimethyl lysine IP from nuclear lysates from tumors treated as in G for 19 days. (I) Representative H&E and IHC images of HT29 colon tumors from the mice treated for 18 days experiment in E. (J) UMAP dot plot of scRNAseq data of pretreated (left) or on treatment (right) samples from patients with *BRAF^V600E^* CRC. Samples are colored by cell type/cluster. TAs are transit amplifying cells and EECs are enteroendocrine cells. (K) Relative differences in cell proportions for each cluster between the pretreatment and on treatment samples. Red dots have an FDR < 0.05 and mean absolute |log2FD| > 0.58 in on treatment vs. pretreatment cells.

To determine how LSD1 inhibition altered response to targeted therapy, we implanted HT29 cells orthotopically into the colons of NSG mice and, following tumor formation, treated the mice with vehicle or BRAFi+EGFRi with or without the LSD1i SP-2577. As demonstrated previously, BRAFi+EGFRi treatment reduced tumor growth and levels of phosphorylated ERK and Ki67 (Fig. 4E, S4C,S4D). The addition of SP-2577 further reduced tumor size and Ki67 staining (Fig. 4E, S4C, S4D). Ex vivo luminescence demonstrated that metastasis to the lung was decreased in the BRAFi+EGFRi treated groups, with a trend to further reduction with the addition of SP- 2577 (Fig. 4F). In tumor tissue collected after 7 days of treatment, the interaction of STAT3 with LSD1 increased in BRAFi+EGFRi relative to vehicle tumors but this increase was attenuated with the addition of SP-2577 treatment (Fig. 4G). We previously demonstrated that LSD1- CoREST2 demethylates STAT3 to promote STAT3 activity (12). Here, in tumor lysates prepared from mice treated for three weeks, levels of demethylated STAT3 decreased in BRAFi+EGFRi relative to vehicle tumors but increased in BRAFi+EGFRi+SP-2577 tumors (Fig. 4H).

Histologically, the addition of SP-2577 treatment reduced the BRAFi+EGFRi-induced increase in intratumoral cystic areas and synaptophysin staining (Fig. 4I, S4E). Collectively, these findings suggest that LSD1 inhibition by SP-2577 treatment enhanced the ability of BRAFi+EGFRi to reduce tumor growth by reducing the targeted therapy-induced interaction of LSD1-CoREST2-STAT3 and subsequent lineage plasticity.

### Targeted therapy enriches for secretory cells in a subset of patients with *BRAF^V600E^* CRC

To explore if targeted therapy induced lineage plasticity in patients, we analyzed publicly available scRNAseq data from a phase 2 clinical trial where patients with refractory metastatic *BRAF^V600E^* CRC were treated with BRAFi (dabrafenib), MEKi (trametinib) and immunotherapy (anti-PD-1, sparatlizumab) (19). Pretreatment and day 15 on-treatment scRNAseq data was available for 23 patients. However, samples from only three patients had enough cells in both samples to proceed with analysis (greater than 200) and one of those samples had no expression of EEC marker genes. Therefore, we integrated the scRNAseq data from the two remaining samples and, based on marker analysis, identified clusters representing several types of secretory cells, including secretory progenitors, EECs, goblet cells and tuft cells (Fig. 4J, S4F, Supplementary Table S2). EECs and goblet cell clusters were the most enriched clusters in the on-treatment relative to the pretreatment samples (Fig. 4K). Interestingly, like some of our in vivo models, *PROX1* expression was highest in the secretory cell populations and the number of cells expressing *PROX1* increased in the on-treatment samples (Fig. S4G). Additionally, the EEC cluster showed the greatest enrichment of genes that were downregulated after LSD1 knockout in the pituitary relative to the other cell populations (Fig. S4H). Altogether, these findings suggest that targeted therapy also enriches for secretory cells populations, including EECs, in patients.

## Discussion

Targeted therapy induced lineage plasticity is an emerging mechanism of therapy resistance that has been most well studied in prostate and lung cancer (3). Here, we demonstrate that *BRAF^V600E^*CRC also undergoes therapy induced lineage plasticity following BRAFi+EGFRi treatment. The neuroendocrine cancers induced by treatment in prostate and lung cancer are highly aggressive, metastatic and therapy resistant, which suggests that lineage plasticity toward a neuroendocrine cancer in *BRAF^V600E^* CRC may contribute to the poor outcomes of patients with this cancer.

As EECs are the neuroendocrine cell of the intestine, it is likely that many pathways contributing to neuroendocrine transformation in prostate and lung cancer are also relevant to *BRAF^V600E^* CRC and vice versa. Additionally, there is existing literature on the role of signaling pathways and other factors in the differentiation of the different types of epithelial cells in the normal intestine that will be informative to therapy induced lineage plasticity in other epithelial cancers. Interestingly, the combination of Wnt and MAPK inhibition in the intestine results in EEC differentiation (13). *BRAF^V600E^* CRC typically has lower Wnt activation than *BRAF* wild type CRC and treatment with BRAFi+EGFRi inhibits MAPK. The resultant state of cell signaling pathway levels may be connected to the enrichment of EECs following BRAFi+EGFRi in *BRAF^V600E^* CRC.

Epigenetic factors play significant roles in the maintenance of stemness and differentiation pathways. Here, we demonstrate that LSD1 promotes therapy-induced enrichment of EECs in *BRAF^V600E^* CRC. In addition to LSD1, other epigenetic factors have been implicated in lineage plasticity in other cancer types and it will be of interest to explore them in future studies in CRC (3,11,20). There are several LSD1 inhibitors in clinical trials (11). However, of the ones we tested, only SP-2577 (seclidemstat) was effective in reducing EEC differentiation. SP-2577 has greater ability to block interactions between LSD1 and other proteins than the other LSD1 inhibitors and has been shown to prevent neuroendocrine differentiation in prostate cancer (21). We demonstrated that SP-2577 attenuates the inhibitor-induced interaction of LSD1 with CoREST2 and STAT3, which we have previously shown is required for EEC differentiation (12). Therefore, it is likely that SP-2577 is effective in our models because of its ability to block LSD1 protein interactions, not through inhibiting LSD1 catalytic activity. Interestingly, our data suggests that SP-2577 does not alter the interaction between LSD1 and CoREST1, which could be followed up on to design a LSD1 inhibitor with more specificity.

Matched pretreatment and on treatment samples from patients with *BRAF^V600E^* CRC are difficult to obtain because the cancers are rarely biopsied during treatment. In this study, we used the only publicly available dataset of paired pre and on-treatment samples from this patient group (19). Similar to our preclinical models, on treatment samples had enrichment of secretory cell types, including EECs, compared to pretreatment samples. However, this analysis had several limitations. The sample size was small because only three of the 23 samples with pre and on treatment data had enough cells in the scRNAseq data to proceed with our analysis. Only two of the three samples contained any level of expression of EEC marker genes, suggesting that EEC enrichment occurs in a subset a patients with *BRAF^V600E^* CRC receiving targeted therapy. Additionally, the patients received BRAFi (dabrafenib), MEKi (trametinib) and immunotherapy (anti-PD-1, sparatlizumab) because established dosing and safety data already existed for this regimen from patients with melanoma (19). Whether or not the standard of care treatment for refractory metastatic *BRAF^V600E^* CRC that consists of treatment with encorafenib (BRAFi) and cetuximab (EGFRi) has the same effect on cell type proportions needs to still be tested (2). Interestingly, MEKi increased expression of marker genes for EECs and goblet cells in *BRAF^V600E^* CRC cell lines, which is consistent with the on-treatment samples from the patient data having enrichment of several secretory cell types, not just EECs.

EECs in the normal colon secrete factors that regulate other epithelial and non-epithelial cells such as immune cells and neurons (7). How EECs enriched in *BRAF^V600E^* CRC following targeted therapy influence the tumor microenvironment will be explored in future studies using our newly developed syngeneic model of *BRAF^V600E^* CRC. The stromal differences observed in tumors from this model in BRAFi+EGFRi versus vehicle treated mice suggest that treatment directly or indirectly through changes in tumor cell type influences stromal content. LSD1 inhibitors have also been used to increase tumor antigen presentation (22) so future work will also explore the effect of LSD1 inhibition on the immune response following BRAFi+EGFRi treatment of *BRAF^V600E^* CRC.

BRAFi+EGFRi treatment has limited effectiveness in patients with *BRAF^V600E^* CRC (2). Therapy-induced enrichment of EECs could contribute to resistance to this therapy regimen. Combining targeted therapy with epigenetic therapy such as an LSD1 inhibitor has the potential to improve patient responses to targeted therapy by blocking lineage plasticity.

## Methods

### Cell line and organoid growth and treatment

All cell lines were cultured and maintained as described in (12). All cell lines were purchased from ATCC. HT29 and NCI-H508 cells were authenticated and last tested for Mycoplasma using the Universal mycoplasma detection kit (ATCC, 30-1012K) in 2023. All cells used in experiments were passaged fewer than 15 times with most being passaged fewer than 10 times. 817829 284-R (817) colon cancer organoids were derived from a patient derived xenograft model obtained from the NCI Patient-Derived Models Repository and passaged and differentiated as described previously (10). Mouse TP KO CRC organoids were generated and passaged as described in the supplementary methods. Encorafenib (MedChemExpress #HY- 15605), gefitinib (MedChemExpress #HY-50895), Stattic (Selleckchem #S7024), and SP-2577 mesylate (Seclidemstat mesylate; MedChemExpress #HY-103713A) were solubilized in DMSO (VWR #97063-136) prior to treatment.

RNA isolation and RT-qPCR was performed as in Ladaika et al (12) and using primers and TaqMan assays listed in Supplementary Table S3. Cq values of genes of interest were normalized to housekeeping gene expression and then to the DMSO control.

### Immunofluorescence

Immunofluorescence for βIII-Tubulin was performed as described in Ladaika et al (12) using antibodies listed in Supplementary Table S4.

### Dose response curves

Dose response curves were performed by performing serial dilutions of media containing encorafenib or gefitinib into media containing DMSO, gefitinib, encorafenib and/or LSD1 inhibitors as indicated in figure legends. Following 72H of treatment, cell viability was assayed using CellTiter-Glo Luminescent Cell Viability Assay (Promega #G7572) per manufacturer’s protocol.

### Co-immunoprecipitation

Co-immunoprecipitations were performed from nuclear lysates as in Ladaika et al (12) using antibodies listed in Supplementary Table S4.

### Orthoptic implantation

All mouse experiments were covered under a protocol approved by the Indiana University Bloomington Animal Care and Use Committee in accordance with the Association for Assessment and Accreditation of Laboratory Animal Care International. Approximately 2.0x10^5 HT29 cells expressing firefly luciferase and TdTomato (HT29 LUC2) or dissociated 817 organoids were combined with Matrigel and injected into the mechanically prolapsed large intestines of 6-week-old female NSG mice. Following tumor formation, mice were treated with BRAFi (encorafenib, 20 mg kg^-1^, daily, oral gavage) and EGFRi (cetuximab, 20 mg kg^-1^, biweekly, i.p.) with or without LSD1i (sp-2577 mesylate, 150 mg kg^-1^, twice daily, oral gavage, as in (21)) for three weeks. Two mice in the BRAFi+EGFRi+SP-2577 reached humane endpoints at 7 days post treatment and two mice from each group were euthanized at this time point for comparison. The remaining three mice per group were treated for a total of 3 weeks. For syngeneic tumors, dissociated TP KO organoids expressing luciferase were combined with Matrigel and injected into the mechanically prolapsed large intestines of 4–6-week-old male and female C57Bl/6 mice. Following tumor formation, mice were treated with BRAFi (encorafenib, 20 mg kg^-1^, daily, oral gavage) and EGFRi (gefitinib, 100 mg kg^-1^, daily, oral gavage) for 21 days.

Tumors were measured and tissue was assayed as indicated in the supplementary methods.

### Immunohistochemistry

Immunohistochemistry was performed as in Ladaika et al (12) using antibodies listed in Supplementary Table S4. Staining percentage was scored blindly using ImageJ for color deconvolution and to determine the percent DAB staining of the total tumor area. Tumor grade was evaluated by standard criteria in human tumors with Grade 1,2 regions defined by >50% glandular features and Grades 3, 4 defined by decreased glandular features and increased solid and cord-like structures. All cases were stained by H&E and scanned by Aperio with digital files annotated to include the entire tumor mass in semi-quantitative method with outlining of all tumor; low grade tumor; and high grade tumor by board certified pathologist (GHostetter) and overall tumor composition based on total tumor area (um2) and the area subset grade 3,4.

### Chromium Single Cell Flex Gene Expression

#### Sample preparation, library preparation and sequencing

For sample preparation, 20-25 mg of fresh mouse tumors (7 vehicle and 8 BRAFi+EGFRi) were processed using the Tissue Fixation and Dissociation for Chromium Fixed RNA Profiling protocol (CG000553, 10X Genomics) and the Chromium Next GEM Single Cell Fixed RNA Sample Preparation Kit (1000414, 10X Genomics). See supplementary methods for additional details. Gene expression libraries were prepared at the Indiana University School of Medicine (IUSM) Center for Medical Genomics using the Chromium Fixed RNA Kit, Mouse Transcriptome (1000497, 10x Genomics), in accordance with the user guide (Chromium Fixed RNA Profiling, CG000527- RevD). Briefly, 8,000 cells per sample were targeted for Gel-Beads-in-Emulsion (GEMs) formation after hybridization and pooling equally. The final library was generated and then sequenced using an Illumina NovaSeq X plus with approximately 2500 million reads.

#### Pre-Processing and QC

Read alignment and gene quantification of mouse singe cell RNA sequencing data was done using the CellRanger multi pipeline (version 8.0.1, 10X Genomics; Pleasanton, CA, USA). The pre-built Cell Ranger mouse reference package (mm10) and mouse reference probe set (Chromium Mouse Transcriptome Probe Set v1.0.1) were used for read alignment. Additional QC and preprocessing steps are in the Supplementary Methods.

#### Subsetting of tumor epithelial cells and visualizing data

To identify tumor epithelial cells in the dataset, cells were clustered using the functions *FindNeighbors* and *FindClusters*. Tumor epithelial cells were distinguished from non-epithelial cells using expression levels of epithelial markers and isolated from the Seurat object using the *subset* function. The resulting Seurat object was then normalized, variable features were identified, and the data was scaled, prior to identifying clusters using the *FindNeighbors and FindClusters* function. Cell clusters were then annotated based on marker analysis. Cell proportion changes were calculated using the *propeller* function from the *speckle* package v1.2.0 (23).

#### Analysis of patient scRNAseq data

Filtered matrices and accompanying metadata obtained from the Broad Institute (see Data Availability Statement) were used to make a Seurat object. Similar pre-processing and quality control steps were taken as indicated for the mouse Flex scRNAseq experiment. Prior to clustering, cells were split into separate Seurat objects based on patient identification and treatment status. These Seurat objects were then integrated together using *FindIntegrationAnchors and IntegrateData.* Samples were then renormalized, scaled, and PCA reduction was performed. Clustering and module scores were generated as described earlier. Significance in cell proportion changes were generated using *scProportion* (10).

## Data availability statement

The scRNAseq datasets are accessible via the NCBI’s Gene Expression Omnibus (GEO) through GEO accession number XXXXX. Patient scRNAseq data analyzed in this study were obtained from https://singlecell.broadinstitute.org, study SCP2079. Only samples from patients that had greater than 200 cells for both pretreatment and on treatment samples were used in the analysis. All other raw data generated in this study are available upon request from the corresponding author.

## Acknowledgements

We would like to thank Drs. Anbarasu Kumaraswamy and Joshi Alumkal for their advice on SP- 2577 mesylate use in vivo. We would also like to thank the Indiana University School of Medicine (IUSM) Center for Medical Genomics core for their assistance with performing the scRNAseq, the IU Bloomington Light Microscopy Imaging Center for maintaining and providing access to imaging equipment and the IUSM Histology Core for embedding and sectioning tissue samples. This work was in part supported by a NIH National Cancer Institute Grant (R01CA286090) [to H.M. O’Hagan], a Research Enhancement Grant [to H. M. O’Hagan] from the Indiana University School of Medicine (IUSM), and pilot funding [to H. M. O’Hagan] from the IU Simon Comprehensive Cancer Center (IUSCCC) Tumor Microenvironment & Metastasis Program and the IUSCCC P30 Support Grant (P30CA082709). Preliminary data was also funded in part by the IUSCCC Joe Ward Fellowship [to C. A. Ladaika]. Additional pilot funding was provided by the NCI SPORE Project, Epigenetic Therapies – New Approaches (P50CA254897) Developmental Research Program [to H. M. O’Hagan]. The content is solely the responsibility of the authors and does not necessarily represent the official views of the NIH or IUSM. C. A. Ladaika was supported by the Doane and Eunice Dahl Wright Fellowship generously provided by Ms. Imogen Dahl.

