## Supplemental Figures S1, S2, S3, and S4 for "LSD1 inhibition attenuates targeted therapy-induced lineage plasticity in *BRAF^V600E^* colorectal cancer"

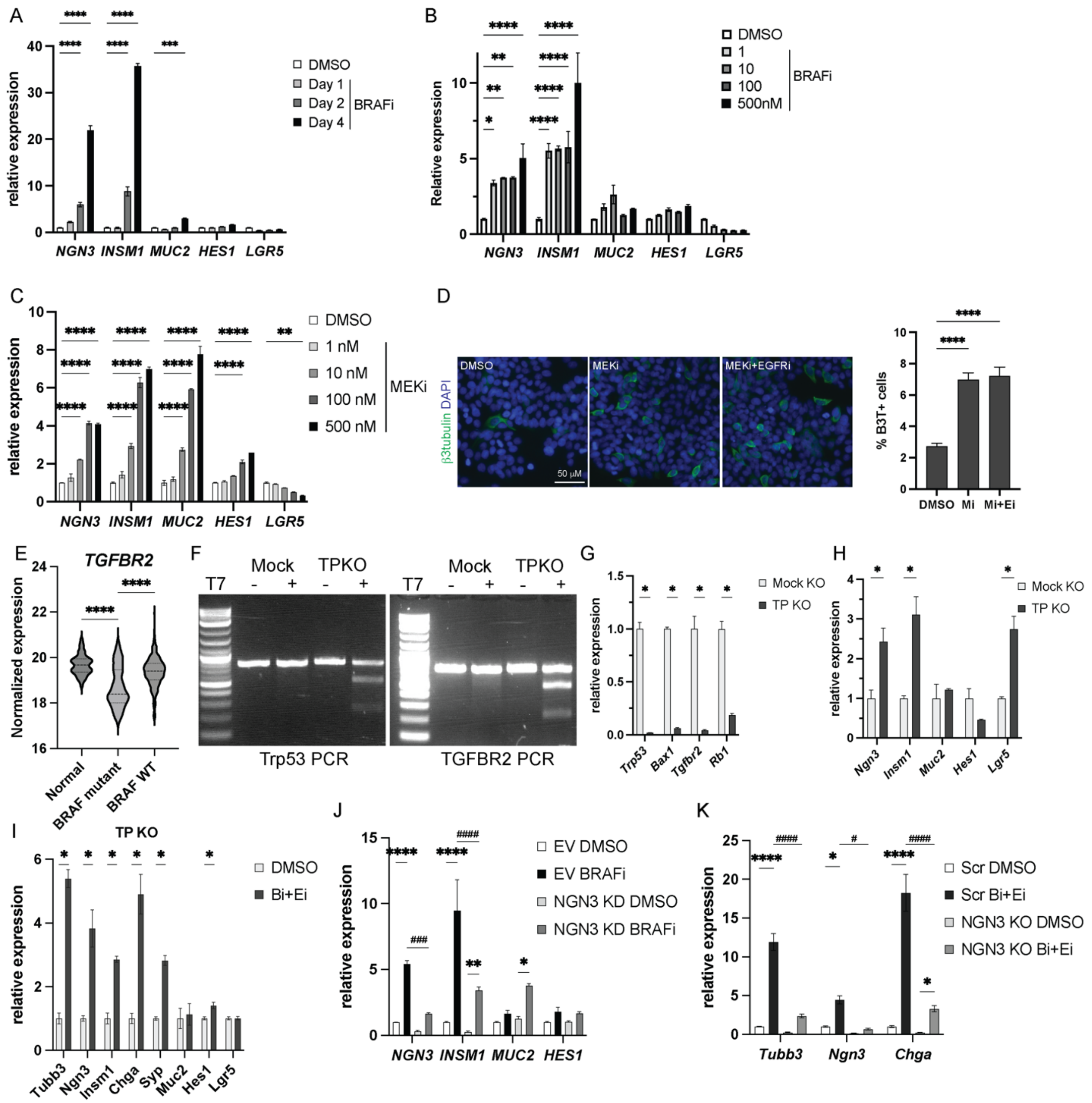


**Supplementary Figure S1. MAPK pathway inhibition enriches for EECs in *BRAF*$^{V600E}$ CRC.** A) Relative gene expression of indicated genes in HT29 cells treated with DMSO or 2.5 nM encorafenib (BRAFi) for the indicated number of days. Gene expression was normalized to the housekeeping gene RhoA and then to the DMSO treated cells. Graph represents mean +/- SEM. N=3. (B) Relative gene expression of the indicated genes in HT29 cells treated with the indicated concentration of encorafenib for 48H. Date presented as in A. (C) Relative gene expression of indicated genes in HT29 cells treated with the indicated concentrations of binimetinib (MEKi) for 48H. Date presented as in A. (D) Immunofluorescence for β3-tubulin (B3T) in HT29 cells treated with 50 nM binimetinib (MEKi, Mi) with or without 500 nM gefitinib (EGFRi, Ei) for 72H. Graph is the %β3-tubulin+ cells of the total number of cells per field. N=3. (E) Normalized expression of *TGFBR2* in the indicated samples from the TCGA colon adenocarcinoma data. (F) T7 endonuclease 1 mismatch detection assay using primers that span the targeted regions for CRISPR knockout (KO) for Trp53 and Tgfbr2 in mock and TP

(Trp53 + Tgfbr2) KO mouse colon CRC organoids. (G) and (H) Relative gene expression of the indicated genes in mock and TP KO mouse organoids. Graph represents mean +/- SEM. N=3. (I) Relative gene expression of the indicated genes in TP KO organoids treated with DMSO or 2.5 nM encorafenib (Bi) and 500 nM gefitinib (Ei) for 72H. Data presented as in A. (J) Relative gene expression of the indicated genes in empty vector (EV) and NGN3 knockdown (KD) HT29 cells treated and presented as in A. (K) Relative gene expression of the indicated genes in scramble KO (Scr) and NGN3 KO TP KO organoids treated as in I and presented as in A. Significance was determined by one-way ANOVA with Tukey pairwise multiple comparison testing (A,B,C,D,E,J,K) and Student's t-test (G,H,I). $^{*}P \leq 0.05$, $^{**} P \leq 0.01$, $^{***} P \leq 0.001$, $^{****} P \leq 0.0001$ relative to DMSO or Mock control. $^{\#}P \leq 0.05$, $^{\#\#\#}P \leq 0.001$, $^{\#\#\#\#}P \leq 0.0001$ relative to EV Bi or Scr Bi+Ei.

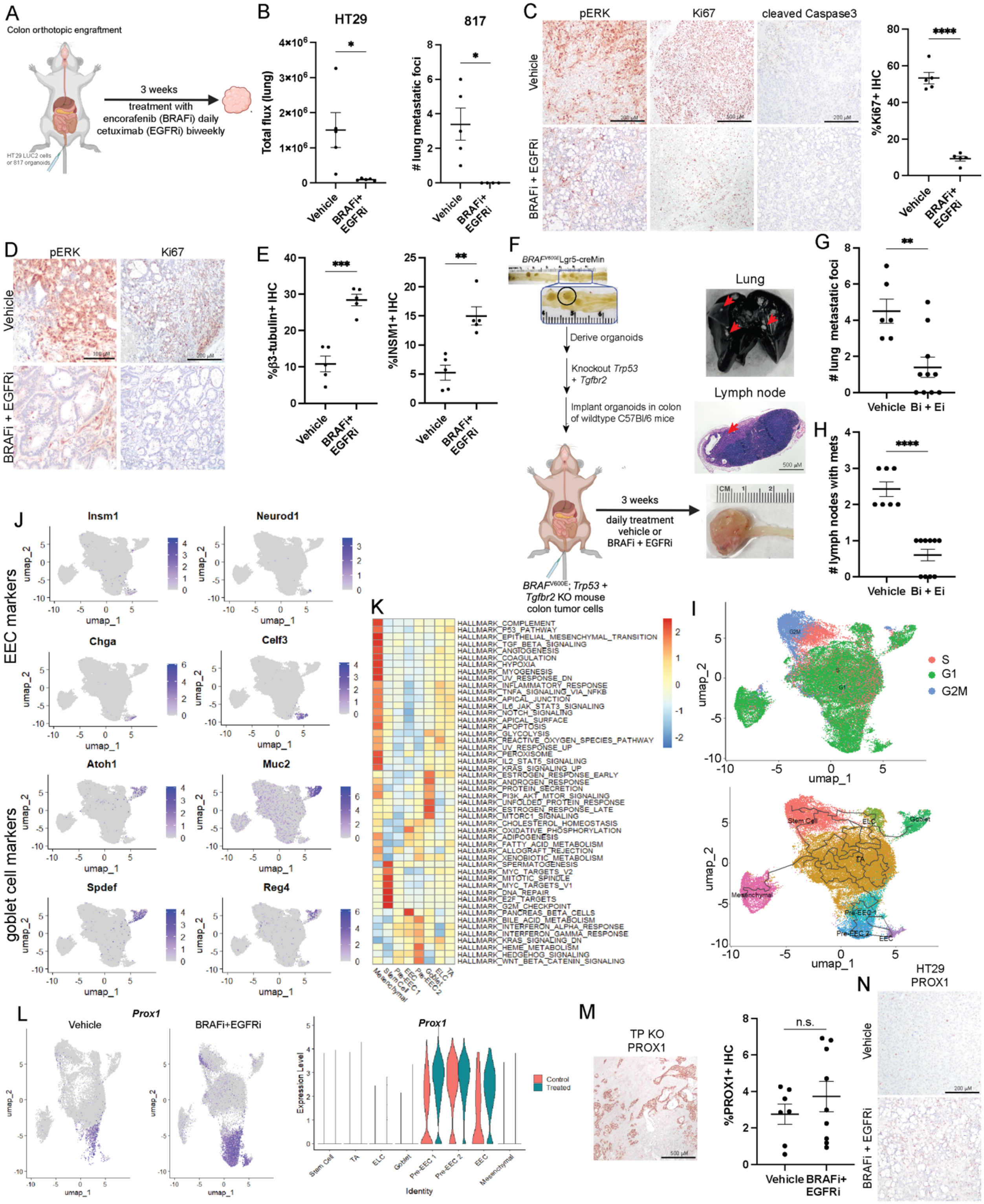

**Supplementary Figure S2. Effect of BRAFi plus EGFRi on tumor cell type composition in *BRAF*$^{V600E}$ CRC.**
(A) Diagram of HT29 and 817 colon orthotopic model and treatment. (B) Total ex vivo flux of lungs from mice with colon orthoptic HT29 tumors treated with vehicle or encorafenib (BRAFi) and cetuximab (EGFRi). Each dot represents the signal from lung from one mouse. Lines

represent mean +/- SEM. (C&D) Representative IHC for indicated proteins in (C) HT29 or (D) 817 tumor sections from mice treated as indicated. (E) Scoring of IHC staining of indicated proteins. Each point represents the scoring for one tumors. Lines represent mean -/+ SEM. (F) Diagram of generation of TP KO organoids and TP KO colon orthoptic model and treatment. (G) Number of metastatic foci in lungs from mice with colon orthotopic TP KO tumors treated with encorafenib (Bi) or gefitinib (Ei) as determined by IVIS imaging. Each dot represents the lung from one mouse. Lines represent mean +/- SEM. (H) Number of lymph nodes with metastases per mouse in mice from F as determined by IVIS imaging. Each dot represents the number of positive lymph nodes in one mouse. Lines represent mean +/- SEM. (I) UMAP dot plots detailing predicted cell cycle phase and trajectory analysis of TP KO scRNAseq samples. (J) UMAP dot plots of normalized expression values of marker genes representative of EEC and goblet cell populations in TP KO scRNAseq samples. (K) Heatmap of enrichment scores for Hallmark gene sets by cell cluster in the TP KO scRNAseq data. (L) UMAP dot plot and violin plot of normalized *Prox1* expression in TP KO scRNAseq data separated by treatment type. (M) Representative image of high PROX1 IHC in TP KO tumors. %PROX1 positive IHC staining as quantified by ImageJ. Each dot represents an individual tumor. Lines represent mean -/+ SEM. (N) Representative PROX1 in HT29 colon orthotopic tumors. Significance was determined by Student's t-test. *P≤ 0.05, ** P ≤ 0.01, *** P ≤ 0.001, **** P ≤ 0.0001, n.s. – not significant.

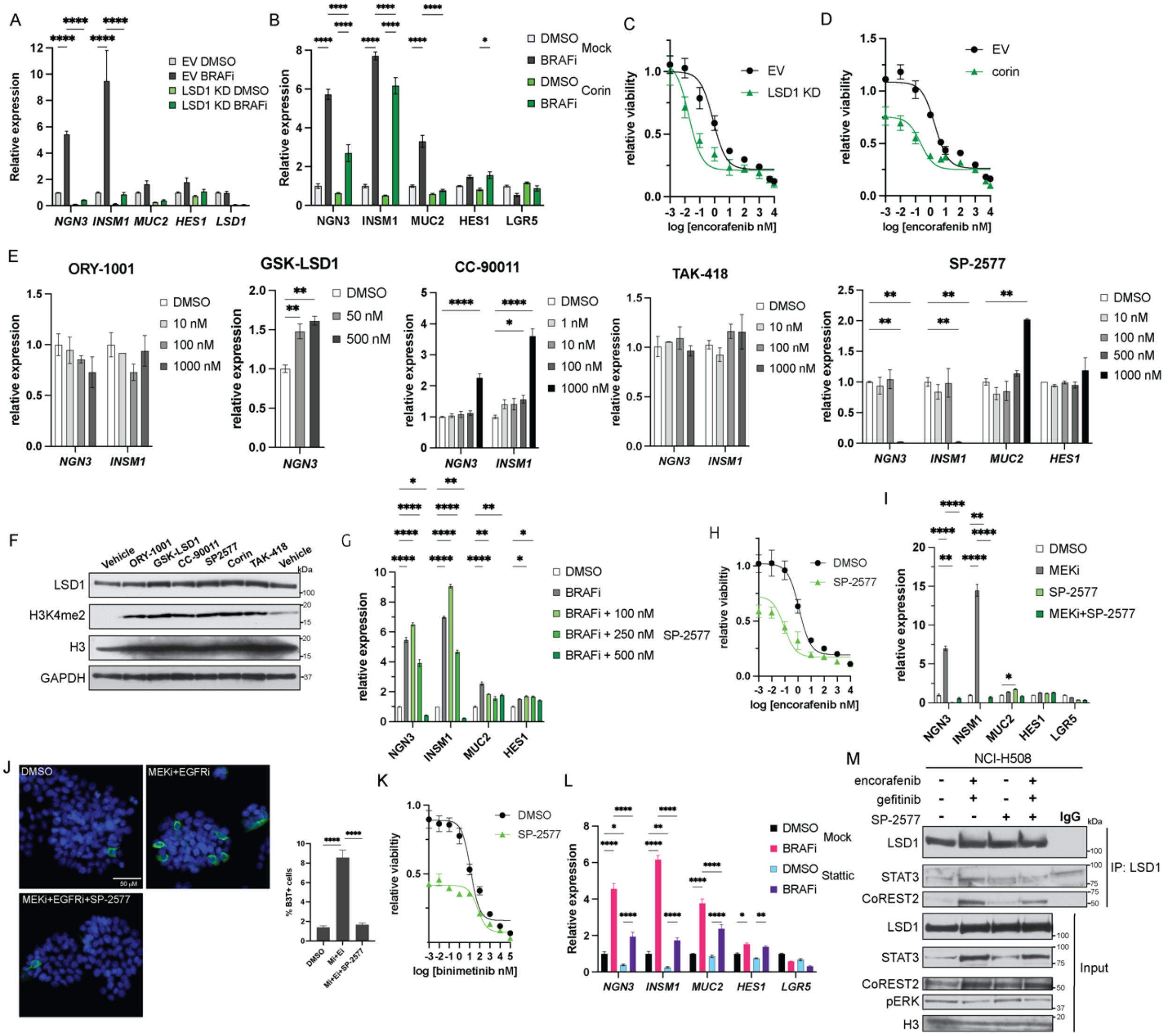


**Supplementary Figure S3. LSD1 depletion blocks MAPK pathway inhibition-induced increase in EECs.** (A) Relative gene expression of indicated genes in empty vector (EV) and LSD1 knockdown (KD) HT29 cells treated with DMSO or 2.5 nM encorafenib (BRAFi) for 48H. Gene expression was normalized to the housekeeping gene RhoA and then to the DMSO treated cells. Graph represents mean +/- SEM. N=3. (B) Relative gene expression of indicated genes in HT29 cells treated with DMSO or 2.5 nM encorafenib with or without 50 nM corin for 48H. Data is normalized and presented as in A. (C) Encorafenib dose response curve of EV and LSD1 HT29 cells treated for 72H. Viability was normalized to cells treated with DMSO only. (D) Encorafenib dose response curve of HT29 cells treated with 50 nM corin for 72H. Viability was normalized to cells treated with DMSO only. (E) Relative gene expression in HT29 cells treated with the indicated concentrations of the indicated LSD1 inhibitors. Graph represents mean +/- SEM. N=3. (F) Western blot of cell lysates prepared from HT29 cells treated with 10 nM of the indicated LSD1 inhibitors. (G) Relative gene expression of indicated genes in HT29 cells treated with DMSO or 2.5 nM encorafenib with or without the indicated concentration of SP-2577 for 48H. Data is normalized and presented as in A. (H) Encorafenib dose response curve of HT29 cells treated with 500 nM SP-2577 for 72H. Viability was normalized to cells treated with DMSO only. (I) Relative gene expression of indicated genes in HT29 cells treated with DMSO or 20 nM binimetinib (MEKi) with or without 500 nM SP-2577 for 48H. Data is normalized and presented as in A. (J) Immunofluorescence for β3-tubulin in HT29 cells treated with DMSO or 20 nM binimetinib (MEKi; Mi) and 500 nM gefitinib (EGFRi, Ei) with or without 500 nM SP-2577for 72H. Graph is the %β3-tubulin+ cells of the total number of cells per field. N=3. (K) Binimetinib dose response curve of HT29 cells treated with 500 nM SP-2577 for 72H. Viability was normalized to cells treated with DMSO only. (L) Relative gene expression of indicated genes in HT29 cells treated with DMSO or 2.5 nM encorafenib (BRAFi) with or without 2 μM Stattic (STAT3i) for 48H. Data is normalized and presented as in A. (M) LSD1 coIP in nuclear lysates prepared from NCI-H508 cells treated with DMSO or 2.5 nM encorafenib plus 250 nM gefitinib with or without 500 nM SP-2577 for 48H. Significance was determined by one-way ANOVA with Tukey pairwise multiple comparison testing. *P≤ 0.05, ** P ≤ 0.01, *** P ≤ 0.001, **** P ≤ 0.0001.

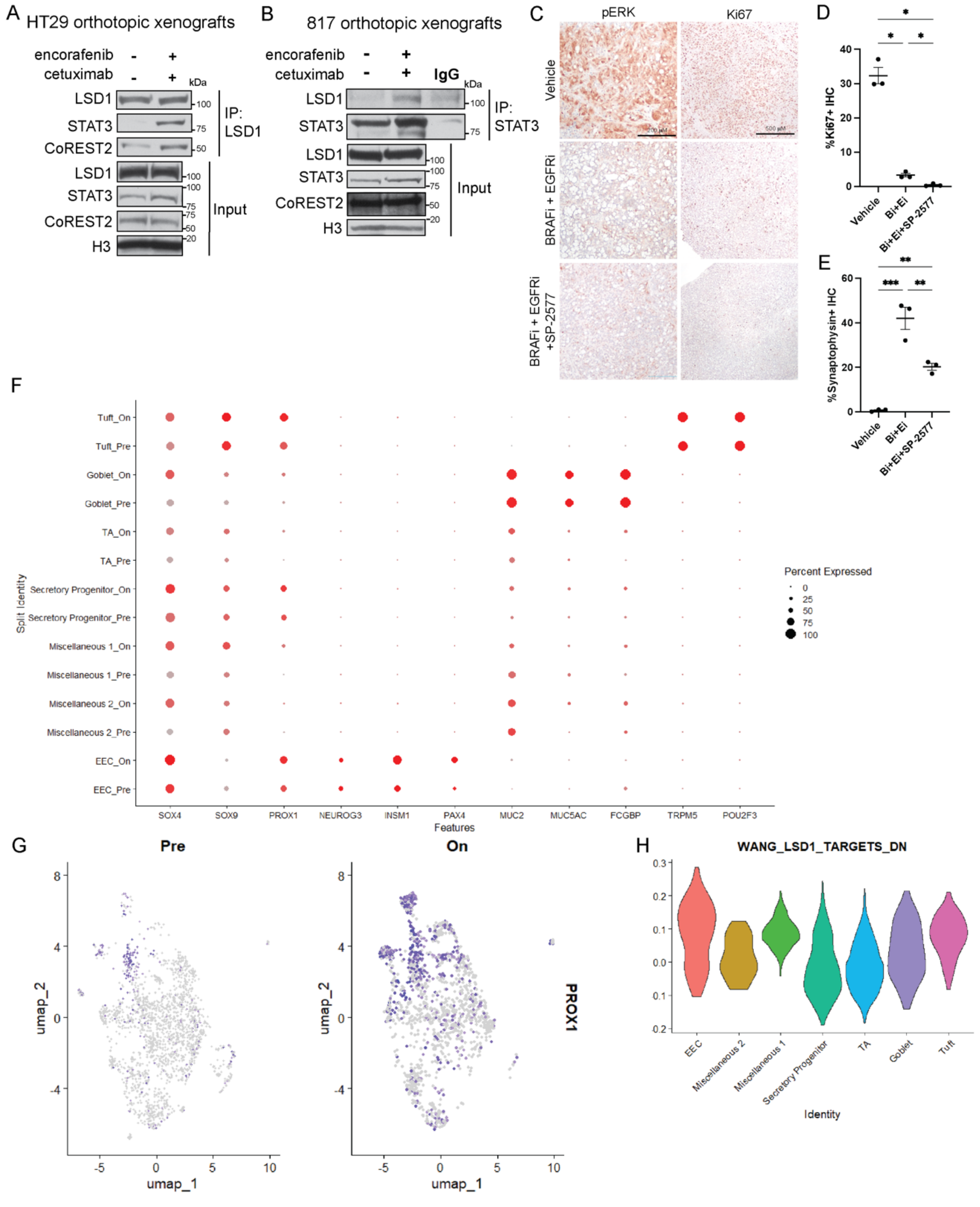


**Supplementary Figure 4. Epithelial cancer cell types present in patient samples of *BRAF*$^{V600E}$ CRC.**

(A) LSD1 coIP in nuclear lysates prepared from HT29 colon orthotopic tumors from vehicle or encorafenib + cetuximab treated NSG mice. The same tumor lysates were also used in the experiment in 4G, so the input blots are the same. (B) STAT3 coIP in nuclear lysates prepared from 817 colon orthotopic tumors from vehicle or encorafenib + cetuximab treated NSG mice. (C) Representative IHC for indicated proteins in HT29 orthotopic tumor sections from mice treated as indicated. (D and E) Scoring of IHC staining of indicated proteins. Each point represents the scoring for one tumor. Lines represent mean -/+ SEM. (F) Dot plot showing marker gene expression in pretreatment and on treatment samples across all annotated cell types. The size of the dot is proportional to the percentage of cells that express a given gene, and the color scale indicates the average scaled gene expression within the specific cell population. (G) UMAP dot plot of normalized *PROX1* expression in samples from patient with *BRAF*$^{V600E}$ CRC scRNAseq data separated by treatment type. (H) Module scores for the Wang LSD1 Targets Down gene set for each cluster in the scRNAseq data from patients with *BRAF*$^{V600E}$ CRC.
